# Optogenetic control of NOTCH1 signalling

**DOI:** 10.1101/2021.09.27.462029

**Authors:** Joanna Kałafut, Jakub Czapiński, Alicja Przybyszewska-Podstawka, Arkadiusz Czerwonka, Cecilia Sahlgren, Adolfo Rivero-Müller

## Abstract

The Notch signalling pathway is a crucial regulator of cell differentiation as well as tissue organisation. Dysregulation of Notch signalling has been linked to the pathogenesis of different diseases. Notch plays a key role in breast cancer progression by controlling the interaction between the tumour cells and the microenvironment as well as by increasing cell motility and invasion. NOTCH1 is a mechanosensitive receptor, where mechanical force is required to activate the proteolytic cleavage and release of the Notch intracellular domain (NICD). Here, we circumvent this step by regulating Notch activity by light. To achieve this, we have engineered a membrane-bound optogenetic NOTCH1 receptor (optoNotch) to control the activation of NOTCH1 intracellular domain (N1ICD) and its downstream transcriptional activities. Using optoNotch we confirm that NOTCH1 activation increases cell proliferation in MCF7 and MDA-MB-468 breast cancer cells in 2D and spheroid 3D cultures. OptoNotch allows fine-tuning ligand-independent regulation of N1ICD to understand the spatiotemporal complexity of Notch signalling.

## Introduction

The Notch signalling pathway is an evolutionarily conserved cell communication system presents in most multicellular organisms. This pathway plays essential roles during cell fate determination during development and tissue homeostasis. Generally, the Notch system controls lateral inhibition and lateral induction, binary cell fate, and boundary formation during embryogenesis [1]. Consequently, Notch signalling plays key roles in vasculature formation [2], osteogenesis [3,4], and plasticity of the nervous systems [5,6]. Cell-to-cell contact between a cell expressing a NOTCH receptor and a ligand-expressing cell is the basis for activation of Notch signalling. NOTCH transmembrane receptors (NOTCH1-4) consist of a large extracellular domain (ECD), a transmembrane domain (TMD), and an intracellular domain (NICD). The ligands, Delta/Serrate/Lag2 family of proteins also have long ECDs followed by a TMD and an ICD [7,8]. NOTCH receptors ECD are cleaved during biosynthesis and non-covalently attached to the rest of the receptor allowing for detachment from the rest of the receptor upon mechanical pulling by any of the ligands. In turn, the unfolding created by the mechanical removal of the large ECD uncovers a cryptic site that is then recognised by proteases (ADAM10, ADAM17 [9], and later on gamma-secretase [10]) causing a series of enzymatic events leading to the cleavage and release of NICD from the membrane. The NICD then translocates to the nucleus where it converts the transcriptional repressor CSL (also known as RBP-J) into a transcriptional activator of Notch-targeted genes [11], such as *HEY* (Hairy/enhancer-of-split related with YRPW motif), *HES* (Hairy/Enhancer of split [E(spl)]) families of genes as well as *MYC* (c-Myc protein), *DTX* (Deltex E3 ubiquitin ligase) and *NRARP* (NOTCH regulated ankyrin repeat protein) [12].

Besides physiological processes, Notch signalling plays roles in progression, migration, invasion, and metastasis in several human malignancies. Additionally, its upregulation is often associated with poor prognosis and drug resistance [13,14]. In breast cancer (BC), Notch signalling is related to the control of proliferation, autophagy, apoptosis [15], cancer cell stemness, and chemosensitivity [16]. Moreover, NOTCH and, in particular, its ligand JAG1 have also been implicated in cluster-cell migration and metastasis of BC [17]. Thus, the monitoring of Notch-related gene expression patterns, as well as the control of Notch signalling, is likely to have therapeutic potential in breast cancers [18–21].

Notch signalling plays a crucial role in controlling basic physiological processes as well as disease development. Yet, Notch signalling is pleiotropic and the outcomes vary, for example depending on the ligand on the sending cell [22]. This has allegedly been to be the result of different modes of activation e.g. short impulse vs long activation patterns [23], as the result of the ligands grip and affinity for the ECD of Notch receptors [24,25]. Therefore, to effectively study spatially and temporally Notch signalling, proper tools are needed.

Optogenetics utilises light-sensitive proteins to stimulate biological processes in an illumination-dependent manner. Optogenetic tools enable non-invasive, flexible, and inexpensive modulation of biological processes such as activation or inactivation of biological pathways [26,27], control of gene expression [28,29], and DNA recombination [30]. Photoactivation has many advantages comparing to traditional methods, such as the use of chemicals or genetic systems. Light control allows avoiding off-target interactions between chemicals and cellular components. Additionally, light can be precisely directed to a single cell, or an area of a cell, and can be carefully controlled in intensity and duration.

Here, we show that an engineered OptoNotch system can regulate NOTCH1 activity with spatiotemporal precision. We apply optoNotch to induce breast cancer cell proliferation and spheroids growth. The fine-tuned regulation of NOTCH1 activity makes this an excellent tool to study the role of Notch signalling in embryogenesis, cancer biology, and drug resistance.

## Results and Discussion

NOTCH1 was re-engineered so the N1ICD is released from a membrane tethered location. To achieve that we adapted the reversible optogenetic system LOVTRAP. This system is based on two proteins, Light-oxygen-voltage-sensing domain 2 (LOV2) and Zdark-1 (Zdk1), that dimerise in the dark but dissociate under blue light stimulation [31]. We anchored LOV2 to the cell membrane, the natural location of inactive NICD, via the Myristoylation-targeting sequence (MTS), while fused Zdk1 to the intracellular domain of the human NOTCH1 (hN1ICD) **(Fig. 1A)**. To facilitate the expression of both proteins in cells, we joined them through a P2A (porcine teschovirus-1 2A) “self-cleaving” sequence so that both could be expressed within one reading frame **(Fig. 1B)**. The P2A is a self-cleaving peptide, which enables the formation of two separate proteins after translation **(Fig. 1C)**. In the dark, LOV2 and Zdk1-NICD are located at the cell membrane. Upon release from the cell membrane, the hN1ICD translocate to the nucleus where it acts as a transcriptional activator together with CSL and MAML **(Fig. 1A)**. We initially tested a construct containing wild-type LOV2^WT^, however, the dissociation of Zdk1 from LOV2 is a reversible reaction, and they rapidly reunite with high affinity in the absence of illumination. This reaction resulted in a very low luciferase signal after photoactivation of the optoNotch system no matter the length of activation (0.05-second pulses for 1, 3, or 12 hours) **(Fig. 1E)**. We then generated the V416L mutation in LOV2 (LOV2^V416L^) which is known to result in a slower regain of affinity for Zdk1 after photo-dissociation [31]. This mutant showed a clear increase in reporter expression upon blue light activation (0.05-second pulses for 1, 3, and 12 hours) by approximately 5-fold as compared to the WT LOV2 **(Fig. 1F)**.

**Figure 1.**
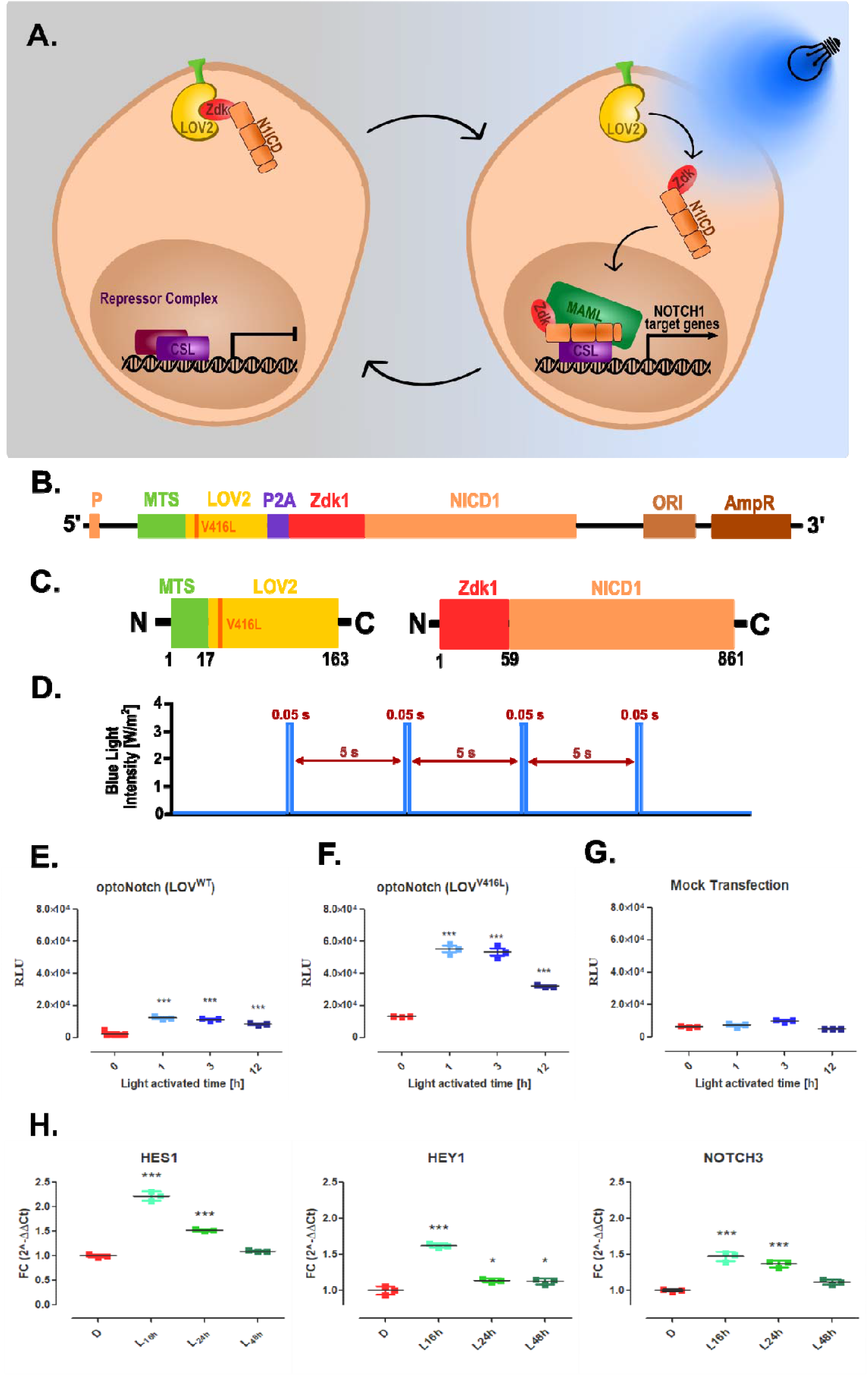
Engineering of photo-activatable Notch. Scheme of the engineered system and optogenetic mode of action used throughout this study. **(A)** Schematic representation of the proposed NICD1 optogenetic control. The LOV2^WT^:Zdk1-NICD1 or LOV2^V416L^:Zdk1-NICD1 is anchored in the cell membrane until photo-activation (456 nm, blue thunder) induces dissociation of the complex, where Zdk1-N1ICD translocates to the nucleus to activate Notch-target genes. **(B)** The vector is designed to express LOV2^WT^ or LOV2^V416L^ and Zdk1-NICD linked by a P2A “self-cleaving” sequence; **(C)** resulting proteins containing LOV2^WT^/^V416L^ (with membrane targeting sequence, MTS) and Zdk1-NICD1. P - CMV promoter, MTS - myristoylation-targeting sequence, LOV2-light-oxygen-voltage-sensing domain 2, P2A-porcine teschovirus-1 2A, Zdk1-Zdark-1, NICD1– human NOTCH1 intracellular domain. **(D)** Schematic representation of blue light activation pattern (pulses) used in optoNotch system **(E-G)** The HEK293T cells expressing either LOV2^WT^-Zdk1-NICD1 or LOV2^V416L^-Zdk1-NICD1 were transfected by the *12xCSL*-Luc reporter and subsequently light activated. The relative luminescence units (RLU) level was measured in LOV2^WT^ **(E)** or LOV2^V416L^-Zdk1-NICD1 **(F)** and mock-transfected cells **(G)** 48 h after activation. The duration of photoactivation of optoNotch (0 h, 1 h, 3 h, and 12 h) does not equate to the increased response. **(H)** The mRNA expression of human *HES1, HEY1,* and *NOTCH3* genes was determined by qPCR (2^−ΔΔCt^) 16, 24, and 48 h after light stimulation of optoNotch-HEK293T cells. The results are presented as fold change (FC) values (mean ± SD) normalized to the human *GAPDH* gene expression. All data were analysed with a one-way ANOVA test and Tukey’s multiple comparisons post-hoc test vs. not light stimulated cells (the time point 0 h and D; dark, respectively). *, p < 0.05; **, p < 0.01; ***, p < 0.001 were considered statistically significant.

We then evaluated the effects of short or longer light activation in downstream reporter activity. We expected that the increased length of photoactivation will also result in higher reporter activity. To our surprise, that was not the case, 1 and 3 h activation resulted in identical reporter values while 12 h light exposure resulted in a reduced luciferase signal **(Fig. 1F)**. Mock transfected cells, but having the reporter system, under identical conditions showed that blue light has no effect reporter expression **(Fig. 1G)**.

To confirm that optoNotch maintains the roles of endogenous N1ICD, we analysed the response of Notch-target genes by qPCR after illumination. For this purpose, *HES1*, *HEY1*, and *NOTCH3* genes were selected, whose expression is known to be dependent on NOTCH1 activity [32,33]. We photoactivated optoNotch-expressing cells using pulses for 3 h and analysed the expression of Notch-targeting genes 16, 24, and 48 h after. This had 2 purposes: first, to show that optoNotch maintains its N1ICD functionality as the naïve N1ICD, and second that its activity is reversed without further photo-activation. All three genes showed increased expression in light-activated cells **(Fig1. H, L samples)** as compared to the same cells kept in the dark **(same figure, D samples)**. The greatest increase of target gene expression was observed 16 hours after light activation. As expected, without further photoactivation of optoNotch the expression of target genes slowly returned to their original levels in about 48 h **(Fig. 1F)**.

### OptoNotch on breast cancer cell proliferation

Since NOTCH1 signalling is a well-known player in breast cancer development [34], we selected two breast cancer cell lines, MCF7, and MDA-MB-468 cells, to assess the functionality of optoNotch. Due to the low efficiency of transient transfection in these cell lines, we generated stable lines expressing the MTS-LOV2^V416L^-P2A-Zdk1-NICD1 complex (oN). To ensure the expression oN, we analysed these cells by immunostaining the N1ICD using flow cytometry. The total content of N1ICD in both of these stable cell lines was substantially higher than in WT cells (where there is some endogenous N1ICD) (**Fig. 2B-C**).

**Figure 2.**
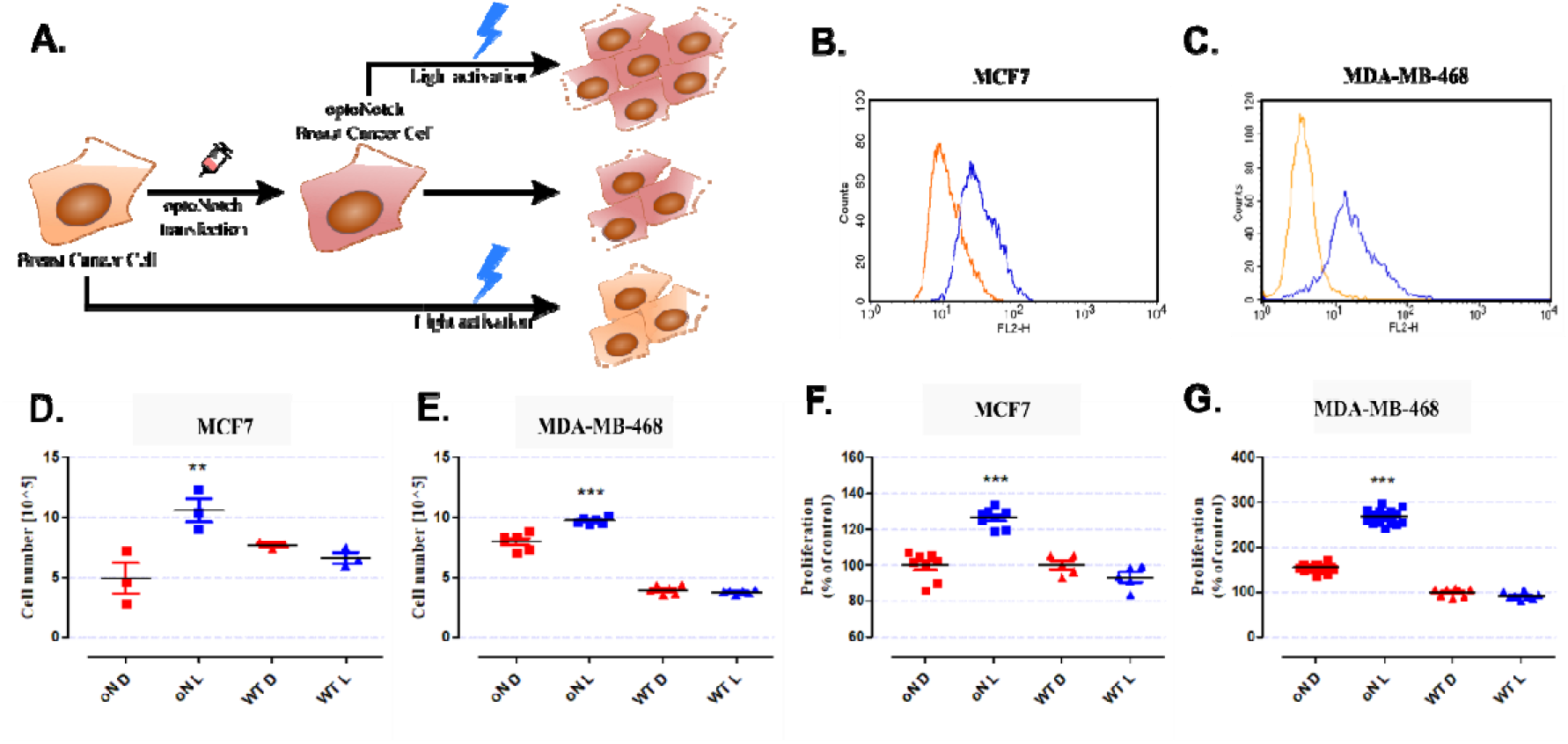
Functional effects of light activation of optoNotch (oN) stable expressing breast cancer cell lines. Schematic representation of the oN on cell proliferation **(A)**. The level of N1ICD was determined by immunostaining using flow cytometry in wild type (orange) and LOV2^V416L^-Zdk1-N1ICD stable expressing (blue) MCF7 **(B)** and MDA-MB-468 **(C)** cell lines. Cells were activated by 0.5 s blue light pulses for 3 hours per day. After 96 h, MCF7 and MDA-MB-468 cells were counted in a TC20™ Automated Cell Counter. Simultaneously, MCF7 and MDA-MB-468 cell proliferation was measured by the MTT assay. The results (mean ± SD) show cell number values **(D, E)** and proliferation (% of control) **(F, G)** of MCF7 and MDA-MB-468 cell lines, respectively. oN D - LOV^2V416L^-Zdk1-NICD1 stable expressing cells kept in the dark; oN L - light-activated LOV2^V416L^-Zdk1-NICD1 stable expressing cells; WT D - wild type cells kept in the dark; WT L – light-activated WT cells. All data were analysed with a one-way ANOVA test and Tukey’s multiple comparisons post-hoc test vs. not light stimulated cells (D; dark). *, p < 0.05; **, p < 0.01; ***, p < 0.001 were considered statistically significant.

We then proceeded to determine the rate of cell growth with and without light activation in oN and wild type (WT) breast cancer cells. The cells were activated by blue light in 0,05/5 s pulses for 1 h per day, as 1 h was the setting resulting in highest reporter gene expression **(Fig 1F)**. After 96 h, the number of cells was counted by using a cell counter. In a parallel experiment, cell proliferation was determined by the MTT assay. Both MCF7 and MDA-MB-468 cell lines in each test showed an increased rate of proliferation after lightactivation (oN L) as compared to the same cells kept in the dark (oN D). WT cells of both lines, whether light-activated (WT L) or not (WT D), maintained an identical proliferation rate between them **(Fig. 2D-G)**.

To confirm the influence of the optoNotch system on breast cancer development under physiological-mimicking conditions, both lines with and without optoNotch were cultured in 3D spheroids using Matrigel. The γ-secretase inhibitor DAPT (10 μM) was used to rule out an effect on the growth of spheroids by endogenous Notch activity. Photo-activation of optoNotch induced faster proliferation - the size of the spheroids increased faster than those in dark (**Fig. 3A and C**), or those without optoNotch whether they were illuminated or not (**Fig. 3B and D**).

**Figure 3.**
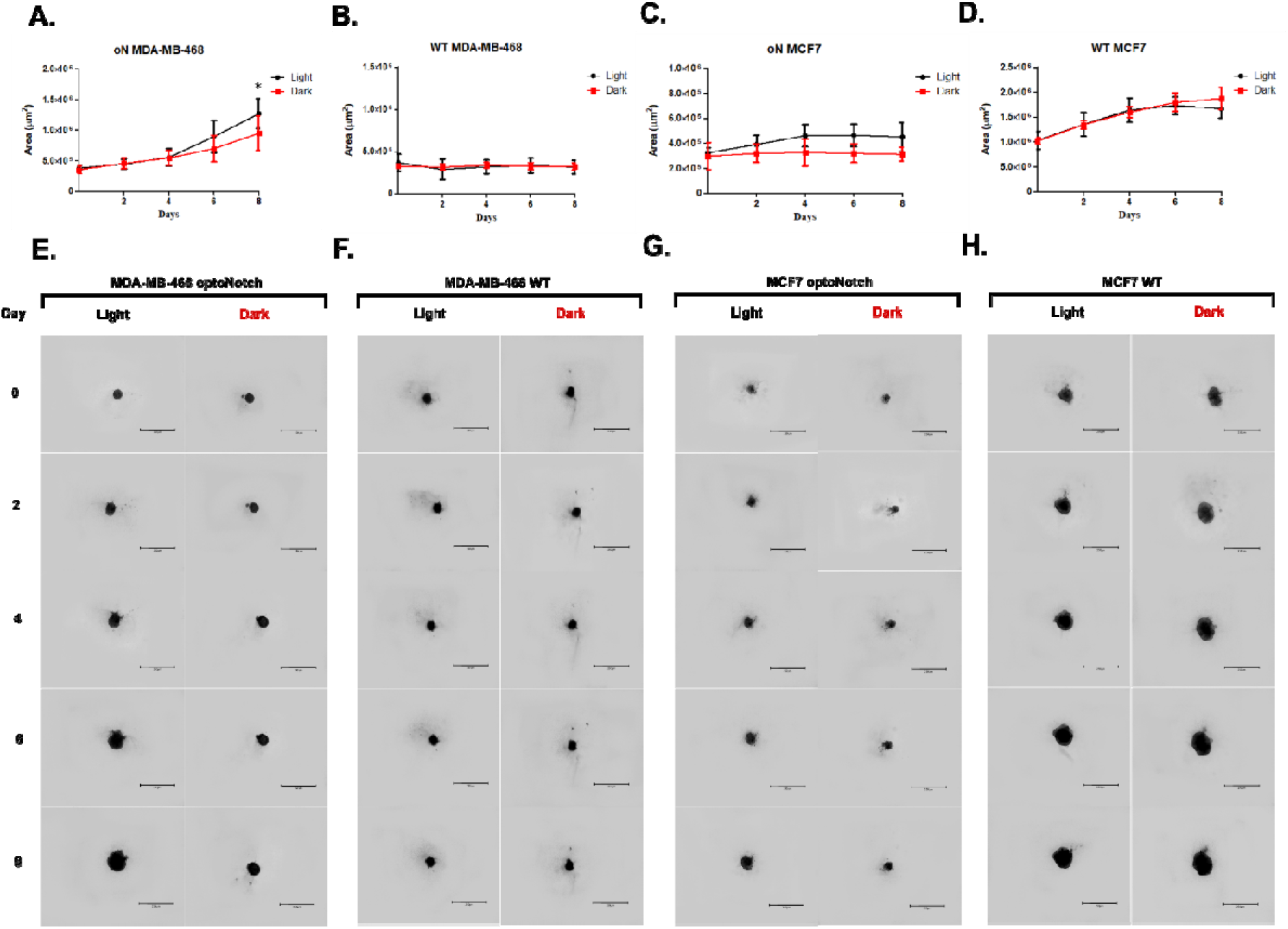
The total size of breast cancer cells spheroids. The LOV2^V416L^-Zdk1-NICD1 mutant and wild type (WT) MCF7 and MDA-MB-468 cells were cultivated in Matrigel with 10 μM DAPT in duplicated 96-well plates. The spheroid (n=7) area was analysed every 48 h for 8 days. **(A)** Stable MDA-MB-468 and **(C)** MCF7 cell lines expressing LOV2^V416L^-Zdk1-NICD1 (oN) and wild type (WT) cells **(B and D, respectively)** are shown. Light (black line) - cells light-activated for 3 hours every day; Dark (red lines) - not light stimulated cells. Panel **(E-H)** representative images of stable LOV2^V416L^-Zdk1-NICD1 and WT breast cancer spheroids were taken every 48h for the duration of the experiment (8 days).

We verified previous research where high NOTCH1 activity fuels breast cancer cell proliferation [35], and we can promote this in the absence of mechanical activation of NOTCH1, as endogenous Notch activity was blocked by a γ-secretase inhibitor (DAPT). Using optoNotch we can modulate receptor activation to discriminate between different modes of NOTCH1 activation, as suggested by differential binding and force used by each of the ligands [36]. The fine-tuned activity of Notch receptors is, likely, the way to fully understand why these receptors often are found to have opposing or variable downstream responses. Indeed, a similar system allowing spatiotemporal control of NOTCH1 activation was recently reported in *Drosophila* [37] where it was used to elucidate transcriptional Notch activity during embryogenesis.

## Materials and Methods

### Materials

KOD-Xtreme hot-start DNA polymerase (Merck Millipore), DreamTaq^™^ Green PCR Master Mix (ThermoFisher Scientific), *DpnI* restriction enzyme (Thermo Fisher Scientific), Gibson Assembly^®^ Master Mix (NEB), Ampicillin (BRAND), Kanamycin (Sigma Aldrich), Spectinomycin (Sigma Aldrich), DNA Clean & Concentrator and Zyppy Plasmid Kits (Zymoresearch), Turbofect^™^ Transfection Reagent (ThermoFisher Scientific), Lipofectamine^®^ 3000 Reagent (Invitrogen), Bright-Glo Luciferase Assay System (Promega), DAPT (Sigma Aldrich), Dulbecco’s Modified Eagle Medium (DMEM/F-12), Penicillin and Streptomycin (Sigma Aldrich), fetal bovine serum (PromoCell) and geneticin G418 (ThermoFisher Scientific). Plasmids pTriEx-NTOM20-LOV2 and pTriEX-mCherry-Zdk1 were a gift from Klaus Hahn (Addgene plasmid # 81009; http://n2t.net/addgene:81009; RRID:Addgene_81009 and Addgene plasmid # 81057; http://n2t.net/addgene:81057; RRID:Addgene_81057) [31]. Plasmid pDONR223_NOTCH1_ICN was a gift from Jesse Boehm, William Hahn & David Root (Addgene plasmid # 82087; http://n2t.net/addgene:82087; RRID:Addgene_82087) [38]. NOTCH1 activity reporters *12xCSL-Luc* has previous been described in [39]. Cell lines HEK293T, MDA-MB-468, and MCF7 were obtained from ATCC. Primary anti-NOTCH1 (D1E11) rabbit monoclonal antibody (Cell Signaling Technology, Cat. Nr: #3608), secondary anti-rabbit antibody conjugated with AlexaFluor 532 (Invitrogen Cat. Nr: #A-11009). All PCR primers were bought from Genomed (Warsaw, Poland).

### Molecular cloning

A construct MTS-LOV2-P2A-Zdk1-NICD1 was generated by Gibson Assembly by combining products from PCR reaction previously carried out with the addition of homology arm on primers. The Light-oxygen-voltage-sensing domain 2 (LOV2) sequence comes from pTriEx-NTOM20-LOV plasmid, Zdark-1 (Zdk1) sequence from pTriEX-mCherry-Zdk1, and human NOTCH1 intracellular domain (NICD1) from pDONR223_NOTCH1_ICN. All PCR reactions were performed by KOD-Xtreme high-fidelity polymerase according to the manufacturer’s protocol. Next, PCR products were digested with *DpnI* restriction endonuclease to remove methylated template plasmids and purified using DNA purification kits. The Gibson Assembly reaction was performed according to the original protocol. Next, the reaction product was transformed into previously prepared electrocompetent *E.coli* bacteria. The resulting bacterial colonies, after selection with kanamycin 50 μg/mL, were subjected to colony PCR and sequence-verified.

Next, the MTS-LOV2-P2A-Zdk1-NICD1 plasmid was used for creating LOV2 mutation (LOV2^V416L^). The sequences of the forward and reverse primers for LOV2^V416L^ are 5’-GAACTTTCTCATTACTGACCCAAGATTGCC-3’ and 5’-GGTCAGTAATGAGAAAGTTCTTCTCAATACGTTC-3’.

### Cell culture and transfection

Human embryonic kidney 293T (HEK293T) cell line and breast cancer cell lines (MDA-MB-468 and MCF7) cell lines were cultured in Dulbecco’s Modified Eagle Medium (DMEM/F-12). All cells were supplemented with 10% FBS, penicillin (100 units/mL)/streptomycin (100 μg/mL) and kept in an incubator at 37°C in an atmosphere of 5% CO_2_.

One day before transfection, HEK293T cells were seeded 5 x 10^4^ cells/well into a 24-well plate and co-transfected MTS-LOV2^V416L^-P2A-Zdk1-NICD1 and 12xCSL-Luc plasmids using Turbofect^™^ transfection reagent following the manufacturer’s protocol. Mock transfection contains GFP as a transfection control, and 12xCSL-Luc for measuring endogenous NICD levels was used as a negative control (Nc). All experiments were done in, at least, triplicate.

Similarly, one day before transfection, MCF7 and MDA-MB-468 cells were seeded 5 x 10^4^ cells/well into a 24-well plate. Medium with 2% FBS was used 24 h before MCF7 transfection to increase transfection efficiency. For transfection MTS-LOV2^V416L^-P2A-Zdk1-NICD1 used Lipofectamine^®^ 3000 Reagent following the manufacturer’s protocol. After 48 hours the cells were selected with geneticin (G418) (0.75 mg/ml) for three weeks to generate stable lines.

### Immunostaining and Flow Cytometry

Detection expression of MTS-LOV2^V416L^-P2A-Zdk1-NICD1 construct into MCF7 and MDA-MB-468 stable cell lines were assessed by measurement of the fluorescent intensity from binding the primary anti-NOTCH1 (D1E11) rabbit monoclonal antibody and secondary antirabbit antibody conjugated with AlexaFluor532. Cells were fixed by resuspending in fixation and permeabilization buffer (BD Pharmingen, Cytofix/Cytoperm solution, cat. # 554722) and incubated for 20 min on ice. Next, cells were washed (BD Perm/Wash buffer, cat. numb. 554723) and centrifuged (500xg, 5 min). MCF7 and MDA-MB-468 stable lines and wild-type cells were incubated with primary anti-NOTCH1 antibody (1 h, 37°C and 5% CO_2_) and subsequently after wash step, labelled with secondary AlexaFluor532-conjugated antibody (1 h, 37°C, and 5 % CO_2_). Part of the cells was incubated only with AlexaFluor532 conjugated antibody.

All immunostainings were performed immediately before the flow cytometry analysis. For Flow Cytometry was performed using a BD FACSCalibur (BD) with CellQuest Pro Version 6.0. software. The fluorescence AlexaFluor532 intensity of individual cells was determined as Counts/FL2-H 2D-dot plots at least 10,000 events were measured within an acquisition rate of 300 events/second, approximately.

### Photoactivation

Transfected HEK293T or stabile breast cancer MCF7 and MDA-MB-468 cell lines were exposed to light. The MAGI-01 Opto-stimulation system (Radiometech) with blue LEDs (456 +/-2 nm) was used for activation in pulses 0.05 s luminous and 5 s breaks by 1, 3, 12 hours in blue light with intensity 3.2 W/m^2^, while others were kept in the dark.

### Luciferase Reporter Assay

At 48 hours after blue light activation, transfected HEK293T cells were lysed following to manufacturer’s protocol. The equal volume of lysates and Bright-Glo Luciferase reagent was transferred to a black microplate well and measured using a microplate luminometer (Tecan Infinite 200 PRO). The results were then statistically analysed.

### Proliferation Assay

MCF7 and MDA-MB-468 stable lines were seeded into two 96-well plates at a density of 3□×□10^4^ cells/mL. For the next two days, one plate was activated blue light pulses by 3 hours per day, while the second plate remained in the dark. 96 h after the last activation, the cells were exposed to 10μL per well of MTT solution (5mg/mL in PBS with ions) for 3 h. After incubation, 100μL per well SDS buffer (10% SDS in 0.01 N HCl) was added to dissolve the crystals. Next, the colour product of the reaction was quantified by measuring absorbance at a 570 nm wavelength using a microplate reader (Tecan Infinite 200 PRO).

### Cell counting

To determine the number of cells, MCF7 and MDA-MB-468 optoNotch stable cells were seeded into two 24-well plates at a density of 5□×□10^4^ cells/mL. For the next two days, one plate was activated blue light pulses by 3 hours per day, while the second plate remained in the dark. 96 h after the last activation, the cells were counted using a TC20^™^ Automated Cell Counter (Bio-Rad). All measurements were made in triplicate.

### Spheroid cultures

MCF7 and MDA-MB-468 stably expressing the optoNotch construct or wild type (WT) controls, were seeded 50μL per well into two 96 Well Round U-Bottom Plates, Sphera Low-Attachment Surface (Thermo Fisher) at a density of 4□×□10^4^ cells/mL. The next day into wells with spheroids was added Matrigel (Corning) dissolved in medium with the addition of 10 μM of DAPT to inhibit endogenous NOTCH signalling. The plates were allowed to polymerize overnight at 37□C. One plate was blue light (456 nm) activated with an intensity of 3.2 W/m^2^ in pulses of 0.05 s every 5s for 3 hours every day during the experiment, while the second plate has remained in the dark. The spheroids area was monitored for 8 days. All spheroids were grown in 7 replicates.

### Microscopy of living cells

Images and area measurements were obtained using an Evos M5000 Imaging System (ThermoFisher Scientific). The photos were taken on the day Matrigel was added (time 0) and then every two days (2, 4, 6, and 8 days). Each spheroid was visualized daily using identical microscope settings.

### RNA extraction and RT-qPCR

Trypsin was used to detach the cells after the experiment. After removing the supernatant, the cells pellets were lysed according to the ExtractMe Total RNA kit (Blirt) manufacturer’s protocol. cDNAs were synthesized using the High-Capacity cDNA Reverse Transcription Kit with the addition of a RNase Inhibitor (Applied Biosystems). All primers used for qPCR were tested for specificity and sensitivity. Glyceraldehyde 3-phosphate dehydrogenase *(GAPDH)* was used as a housekeeping gene. PCR reactions were performed with PowerUp SYBR Green Master Mix (Applied Biosystems) through the LightCycler^®^ 480 II instrument (Roche) in triplicates on 96-well plates. The number of cycles needed to reach a specific threshold of detection (CT) was used to calculate relative quantification (RQ). Relative mRNA expression was calculated using the delta CT subtraction and normalized to the expression of *GAPDH*.

**Table 1.**
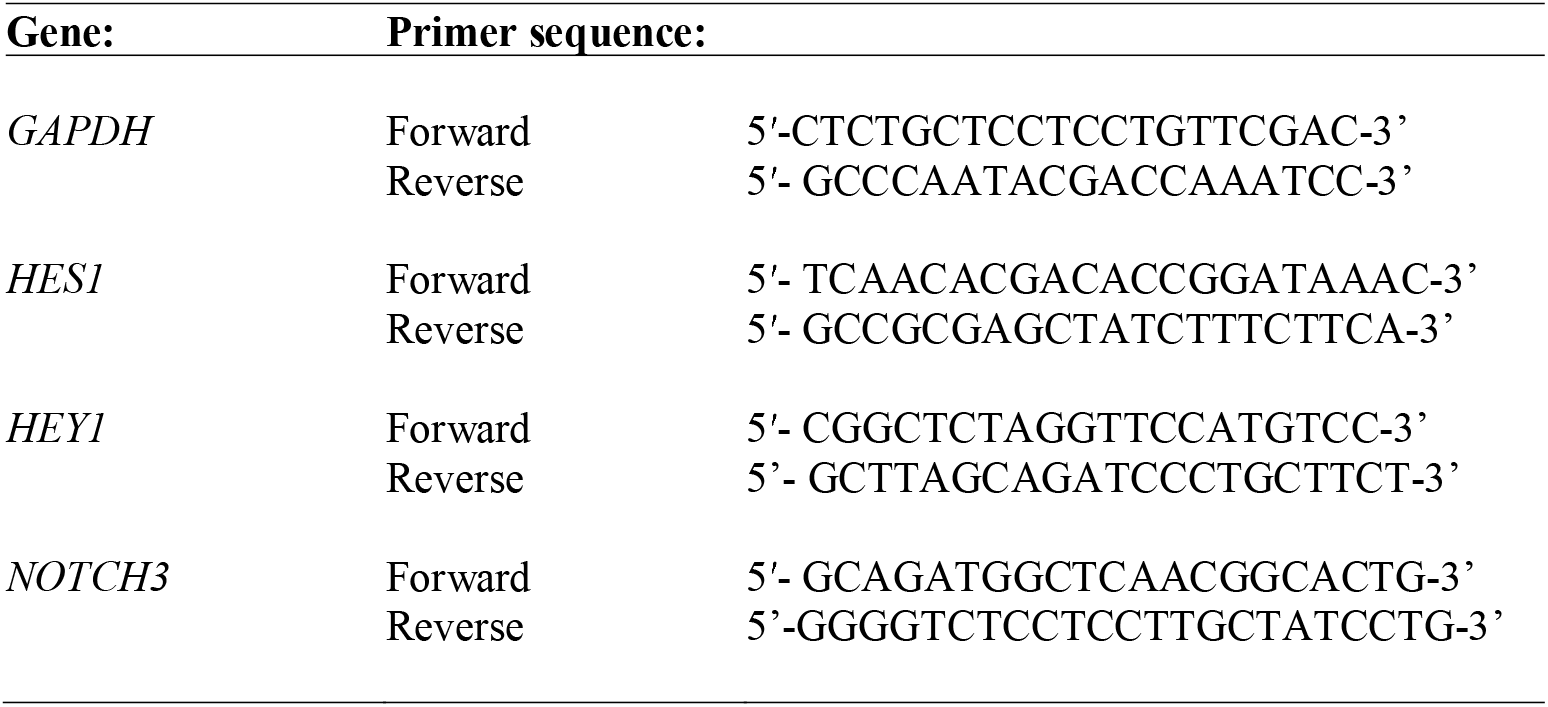
Primers used for qRT-PCR

### Statistical Analyses

Significance among luminescence readings was assessed by one-way ANOVA followed by Tukey’s *post-hoc* test. Statistical analyses of all samples were performed using GraphPad Prism 8.0 (GraphPad Software Inc., California, U.S.A). ANOVA with Tukey post hoc test and column statistics were used for comparisons (*, p < 0.05; **, p < 0.01; ***, p < 0.001 was considered statistically significant). All tests were performed in the triplicates, at least.

## Authors Contribution statement

**JK** carried out the plasmid design, all experiments and the statistical analysis. **APP** and **JK** performed the 3D spheroid experiments. **ACz** assisted with the flow cytometry, qPCR, MTT assay and cell counting experiments. **JK** and **ACz** wrote the manuscript draft. **JK, JCz, CS** and **AR-M** designed the experimental section of this research work. **AR-M** conceptualised and supervised the work. All authors have written and approved the final manuscript.

## Acknowledgements

The work was supported by grants by the Polish National Science Centre (NCN): NZ1/01777 and NZ4/02364.

